# *Staphylococcus aureus* Cas9 is a multiple-turnover enzyme

**DOI:** 10.1101/340042

**Authors:** Paul Yourik, Ryan T. Fuchs, Megumu Mabuchi, Jennifer L. Curcuru, G. Brett Robb

## Abstract

Cas9 nuclease is the key effector of type II CRISPR adaptive immune systems found in bacteria. The nuclease can be programmed by a single guide RNA (sgRNA) to cleave DNA in a sequence-specific manner. This property has led to its widespread adoption as a genome editing tool in research laboratories and holds great promise for biotechnological and therapeutic applications. The general mechanistic features of catalysis by Cas9 homologs are comparable; however, a high degree of diversity exists among the protein sequences, which may result in subtle mechanistic differences. *S. aureus* (SauCas9) and especially *S. pyogenes* (SpyCas9) are among the best-characterized Cas9 proteins and share about 17% sequence identity. A notable feature of SpyCas9 is an extremely slow rate of reaction turnover, which is thought to limit the amount of substrate DNA cleavage. Using *in vitro* biochemistry and enzyme kinetics we directly compare SpyCas9 and SauCas9 activities. Here, we report that in contrast to SpyCas9, SauCas9 is a multiple-turnover enzyme, which to our knowledge is the first report of such activity in a Cas9 homolog. We also show that DNA cleaved with SauCas9 does not undergo any detectable single-stranded degradation after the initial double-stranded break observed previously with SpyCas9, thus providing new insights and considerations for future design of CRISPR/Cas9-based applications.

## INTRODUCTION

Clustered regularly interspaced short palindromic repeats (CRISPR) and CRISPR-associated proteins (Cas) constitute sequence-based adaptive immunity in bacteria and archaea. Our understanding of CRISPR/Cas systems is progressing rapidly, in part driven by the excitement about adapting them for use in research, biotechnology, and human therapy (Chen and Doudna, 2017; Garcia-Doval and Jinek, 2017; Hille et al., 2018; Karvelis et al., 2017; Koonin et al., 2017; Mir et al., 2018; Shmakov et al., 2017; Wang et al., 2016). Of the diverse families of Cas nucleases, *Streptococcus pyogenes* Cas9 (SpyCas9) is the best-characterized and most widely used. SpyCas9 is a monomeric protein that can be programmed with a single guide RNA (sgRNA) to induce sequence-specific double-stranded (ds) breaks in DNA (Jinek et al., 2012). *Staphylococcus aureus* Cas9 (SauCas9) is a less well characterized homolog. The proteins share 17% sequence identity as well as structural and mechanistic parallels (Nishimasu et al., 2015; Ran et al., 2015).

High resolution structures and biochemical studies demonstrated that both SpyCas9 and SauCas9 bind sgRNA by interacting with the 3’-stem loops, which induces a conformational change in the protein (Jiang et al., 2015; Jinek et al., 2014; Mekler et al., 2016; Nishimasu et al., 2015). The Cas9-sgRNA ribonucleoprotein (RNP) complex rapidly screens DNA in search of the protospacer adjacent motif (PAM). Following PAM recognition, the RNP attempts to base pair the sgRNA’s ~20-nucleotide 5’-terminal targeting sequence with the DNA in a 3’-to 5’-direction with respect to the sgRNA. If the DNA sequence adjacent to the PAM is complementary to the sgRNA an RNA:DNA duplex is formed – displacing one of the DNA strands – resulting in an R-loop (Gasiunas et al., 2012; Gong et al., 2018; Jiang and Doudna, 2017; Sternberg et al., 2014; Szczelkun et al., 2014; Zeng et al., 2017). SpyCas9 recognizes a 5’-NGG PAM (Anders et al., 2014; Mojica et al., 2009; Sternberg et al., 2014) motif while SauCas9 recognizes 5’-NNGRRT (Friedland et al., 2015; Nishimasu et al., 2015; Xie et al., 2018). Successful R-loop formation – contingent on perfect (or near-perfect) sgRNA:target DNA match – facilitates DNA cleavage marked by the RuvC domain and HNH domains cleaving the PAM-containing and the non PAM-containing DNA strands, respectively (Gasiunas et al., 2012; Nishimasu et al., 2014; Sternberg et al., 2014). Work *in vivo* and *in vitro,* including single molecule and bulk kinetic experiments showed that upon DNA cleavage SpyCas9 remains bound to the DNA, resulting in extremely slow product release, which ultimately inhibits enzymatic turnover (Gong et al., 2018; Jones et al., 2017; Raper et al., 2018; Richardson et al., 2016; Sternberg et al., 2014). Furthermore, a recent report demonstrates that SpyCas9 modestly degrades cleaved DNA products (Stephenson et al., 2018).

Here, using biochemistry and enzyme kinetics we compared the DNA cleavage activity of SpyCas9 and SauCas9 RNPs, *in vitro,* on a 110 nucleotide-long dsDNA containing a PAM sequence that is recognized by both homologs. Our data suggest that both homologs form highly stable RNPs and in contrast to SpyCas9, which cleaves a stoichiometric amount of DNA, SauCas9 is a multiple turnover enzyme. To our knowledge, this is the first report of such activity among Cas9 homologs. Furthermore, in contrast to SpyCas9, SauCas9 did not have any detectable additional nuclease activity on cleaved DNA products, yielding homogeneous products. Our findings illuminate distinct differences between two Cas9 homologs – both of which are widely used in various biotechnological and therapeutic applications – and add important insights for future development of CRISPR/Cas9 technologies.

## RESULTS

### *S. aureus* and *S. pyogenes* Cas9 bind sgRNAs with comparable affinities and form active, sgRNA-dependent complexes

CRISPR RNA (crRNA) and trans-activating crRNA (tracrRNA) together program Cas9-catalyzed endonucleolytic cleavage of DNA. The two RNAs can be bridged by a GAAA tetraloop, forming a single guide RNA (sgRNA) (Jinek et al., 2012), thus reducing the number of components and facilitating DNA targeting applications. The 3’-end harbors three stem loops in the *S. pyogenes* and two in the *S. aureus* sgRNAs (Nishimasu et al., 2014; Ran et al., 2015), which are thought to be recognized by the protein and are critical for forming an active Cas9-sgRNA RNP. Changing the 5’-proximal ~20-nucleotide sequence is enough to direct Cas9 to a specific site in the substrate DNA. However, not all DNA targets are cleaved with equal efficiency and specificity (Bisaria et al., 2017; Doench et al., 2014). sgRNA stem loops are critical for recognition and binding Cas9 (Briner et al., 2014; Mekler et al., 2016) and it is thought that one contributing reason for differences in cleavage efficiencies is the degree of 5’-end base pairing with the 3’-region of the sgRNA (Thyme et al., 2016; Xu et al., 2017), causing a disruption of the stem loops recognized by Cas9.

We designed SpyCas9 and SauCas9 sgRNAs (Table 1) to have a 20 nucleotide-long targeting sequence (Jinek et al., 2012; Ran et al., 2015) while minimizing the degree of base pairing between the 5’- and 3’-ends – approximated by quick structure prediction algorithms such as mfold (Zuker, 2003) – and measured the binding affinity *(K_D_)* for the respective Cas9 homologs (Fig. 1A). Briefly, sgRNAs were transcribed *in vitro* with T7 RNA Polymerase, purified on a denaturing acrylamide gel, and labeled with a 3’-Cytidine-5 (Cy5) fluorophore. All reactions were carried out at room temperature (22°C) in New England Biolabs Buffer 3.1 (Methods). Cas9 protein was titrated in the presence of 10 nM 3’-Cy5-labeled sgRNA and fraction bound was calculated from the change in fluorescence anisotropy over an increasing concentration of Cas9 protein (Fig. 1A). The data were fit with a quadratic binding equation (Methods), which resulted in comparable *K_D_* values for Spy- and SauCas9 of 21 ± 1 nM and 30 ± 10 nM, respectively (Fersht, 1999; Pollard, 2010).

**Figure 1.**
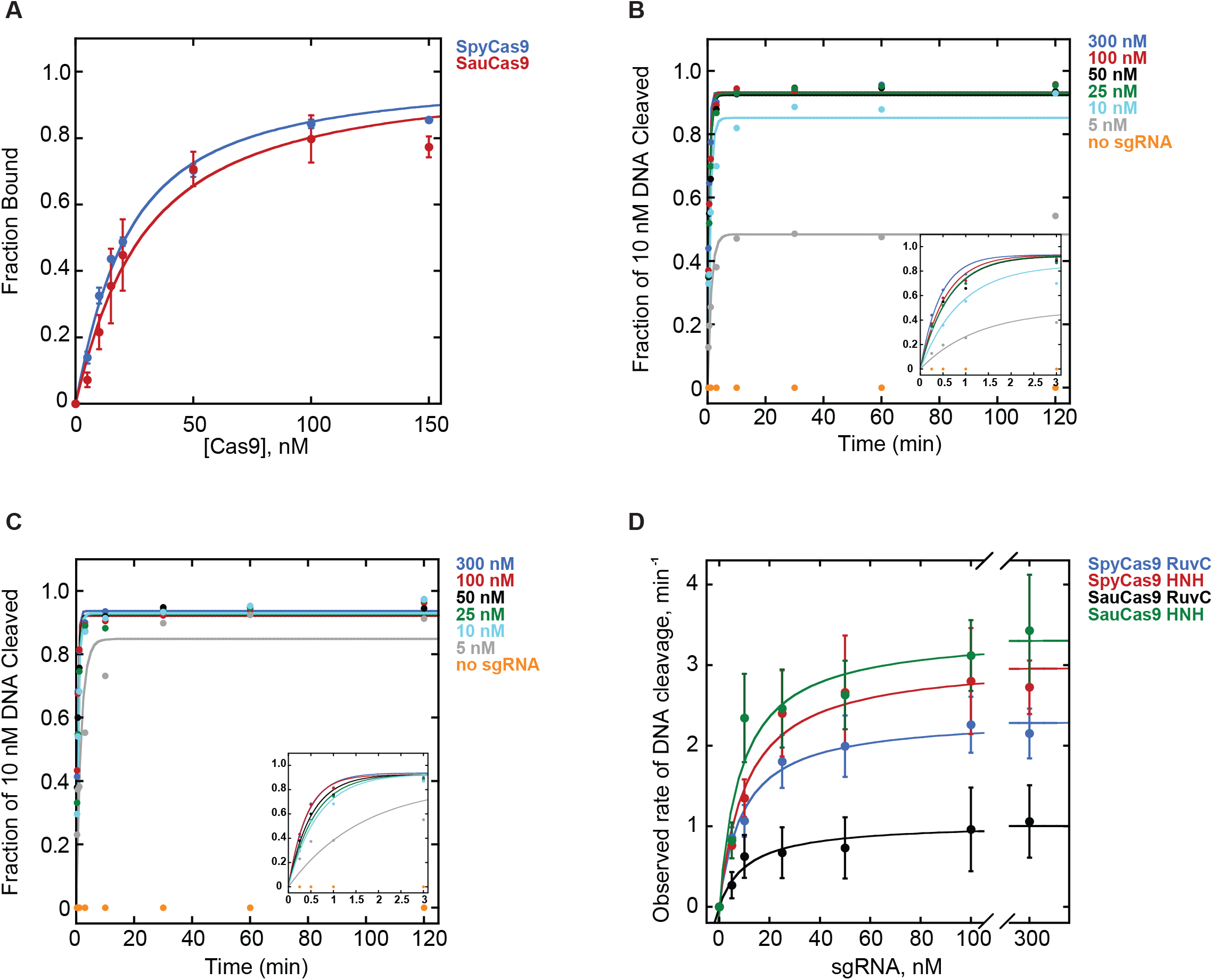
*S. pyogenes* and *S. aureus* Cas9 bind their respective single guide RNAs with comparable affinities and form active RNPs. **A.** Fraction bound of Cy5-labeled sgRNA versus unlabeled SpyCas9 (blue) or SauCas9 (red), measured by fluorescence anisotropy. The *Kd* values for SpyCas9 and SauCas9 were 21 ± 1 nM and 30 ± 10 nM, respectively. **B-C.** Representative plots showing observed rates *(k_obs_)* of HNH domain-catalyzed hydrolysis of 10 nM 110mer DNA by 25 nM (B) SpyCas9 and (C) SauCas9 in the presence of 300 nM (blue), 100 nM (red), 50 nM (black), 25 nM (green), 10 nM (cyan), 5 nM (grey), or no sgRNA (orange). Insets feature early time points of the respective plots for clarity. **D.** Dependence of *k_obs_* on sgRNA concentration for SpyCas9 and SauCas9. Blue – SpyCas9 RuvC domain-catalyzed cleavage: *k_max_* = 2.3 ± 0.4 min^−1^, *K_1/2_* = 8 ± 1 nM; red – SpyCas9 HNH domain-catalyzed cleavage: *k_max_* = 3.0 ± 0.6 min^−1^, *K_1/2_* = 9 ± 1 nM; black – SauCas9 RuvC domain-catalyzed cleavage: *k_max_* = 1.0 ± 0.5 min^−1^, *K_1/2_* = 10 ± 1 nM; green – SauCas9 HNH domain-catalyzed cleavage: *k_max_* = 3.3 ± 0.5 min^−1^, *K_1/2_* = 8 ± 2 nM. All values are reported as mean ± average deviation.

**Table 1.**
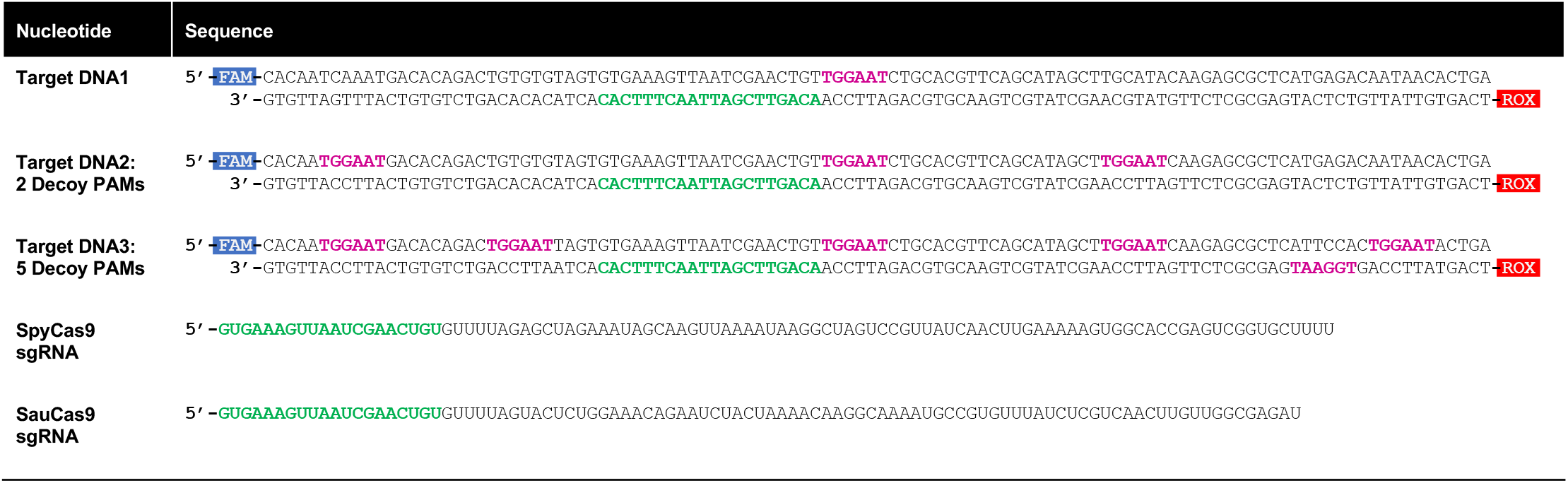
Sequences of target DNAs and sgRNAs used in the study. Target 110mer DNAs were labeled with 5’-FAM (blue box) and 5’-ROX (red box) fluorophores on the forward (PAM-containing) and the reverse (non-PAM) strands, respectively. The PAM region (TGG for SpyCas9, TGGAAT for SauCas9) is in fuchsia. The spacer region, is in green within the DNAs and the sgRNAs.

Having established the affinities with which Spy- and SauCas9 homologs bind their respective sgRNAs, we measured the dependence of the rate and extent of DNA cleavage on the sgRNA concentration. sgRNAs were refolded by heating to 65°C and cooling to 4°C at 0.1°C/sec in a thermocycler. 25 nM Spy- or SauCas9 was preincubated in the presence of a variable concentration of sgRNA, ranging from 0 to 300 nM, for 15 min and reactions were initiated by addition of 10 nM 110mer double-stranded DNA1 (Table 1) harboring a 20-nucleotide target sequence that was a perfect complement to the sgRNA as well as a TGGAAT PAM, which fulfils both Spy (NGG) and Sau (NNGRRT) PAM requirements (Friedland et al., 2015; Mojica et al., 2009). The DNA did not have any other occurrences of either the target sequence or the PAMs (Table 1, DNA1) and was labeled with a 5’-FAM fluorophore on the PAM-containing strand (cleaved by the RuvC domain) and a 5’-ROX fluorophore on the non-PAM strand (cleaved by the HNH domain). Reaction aliquots were quenched with a final concentration of 50 mM EDTA, 1% SDS, and 0.1 units/μL of Proteinase K and analyzed by capillary electrophoresis (CE) (Greenough et al., 2016). The fraction of DNA cleaved was plotted versus time, and fit with a single exponential equation describing the observed rate *(k_obs_)* and extent of DNA cleavage in the presence of a particular concentration of sgRNA (Fig. 1B,C) (Methods). Both Spy- and SauCas9 achieved ≥ 93% extent of cleavage of the DNA when the concentration of the sgRNA was at least stoichiometric to the 110mer target DNA. In contrast, < 2% cleavage was observed in the absence of sgRNA (Fig. 1B,C). *k_obs_* were plotted against the concentration of sgRNA (Fig. 1D) and fit with a hyperbolic equation (Methods), giving the maximal rate of cleavage *(k_max_),* when the reaction is not limited by sgRNA concentration, and the concentration of sgRNA required to achieve the half-maximal rate of DNA cleavage *(K_1/2_).* The *k_max_* for the cleavage of the non-PAM DNA strand by the HNH domain was similar for SpyCas9 and SauCas9 (3.0 ± 0.6 min^−1^ and 3.3 ± 0.5 min^−1^, respectively). The *k_max_* for the cleavage of the PAM-containing strand by the RuvC domain was modestly slower for SpyCas9 (2.3 ± 0.4 min^−1^) consistent with previous reports (Gong et al., 2018; Raper et al., 2018), and approximately 3-fold slower for Sau (1.0 ± 0.5 min^−1^). The *K_1/2_* values for both Spy- and SauCas9 – measured for either PAM-containing or non-PAM strand cleavage – were between 8 nM and 10 nM (Fig. 1D), consistent with an efficient interaction between the Cas9 protein and sgRNA (Fig. 1A). It did not escape our attention that SpyCas9 achieved ~50% reaction extent (~5 nM DNA cleaved) in the presence of substoichiometric (5 nM) sgRNA, while SauCas9 achieved ~95% reaction extent under the same conditions (Fig. 1B,C).

### *S. aureus* but not *S. pyogenes* Cas9 is a multiple turnover enzyme *in vitro*

A prominent feature of SpyCas9 is an extremely slow rate of turnover. Recent work *in vitro* and *in vivo* (Gong et al., 2018; Jones et al., 2017; Raper et al., 2018; Sternberg et al., 2014) demonstrated that the Cas9·sgRNA RNP rapidly finds and cleaves the target DNA sequence but does not dissociate from the cleaved DNA. In contrast, another study involving SpyCas9 suggests that cleaved DNA strands may be released from the post-cleavage complex (Richardson et al., 2016); however, Cas9 is widely accepted to be a single turnover enzyme expected to cleave substrate DNA in approximately 1:1 stoichiometry with active RNP complexes. We performed experiments designed to replicate the previous observations of others by preincubating 25 nM Spy- or SauCas9 and 100 nM of respective sgRNA for 15 min and then added a 10-fold excess of 110mer DNA1 (250 nM) (Table 1), quenched reaction aliquots over a course of 24 hr, and analyzed the reactions by CE as described above. As expected, within less than 5 min of initiating the reaction containing SpyCas9 there was ~25 nM of cleaved product, which increased to 33 ± 5 nM over 24 hr, suggesting that SpyCas9 is nearly 100% active in the presence of saturating sgRNA but there is a very low degree of turnover (Fig. 2A). Strikingly, after 24 hr, SauCas9 resulted in 150 ± 20 nM cleaved DNA, suggesting that the enzyme turns over significantly faster than the *S. pyogenes* homolog. For both SpyCas9 and SauCas9 the reaction was described well by a single exponential followed by a linear (i.e., steady state) phase equation (Methods). The burst kinetics for SpyCas9 were too rapid to be resolved with manual quenching; however, the estimated burst amplitude was consistent with the SpyCas9 concentration of 25 nM (Fig. 2A). It is unlikely that the equation fit to the SauCas9 data recapitulates a true burst because the amplitude is nearly 4-fold higher than the SauCas9 concentration in the reaction (Fig. 2A). Interestingly, the linear phase of the reaction for SauCas9 was 1.6×10^−3^ ± 5×10^−4^ min^−1^, which is 7-fold faster than measured for SpyCas9 (Fig. 2A,D). Preincubating SpyCas9 or SauCas9 RNPs for 24 hr at reaction conditions prior to addition of substrate DNA1 resulted in nearly identical results, strongly suggesting that the RNP is stable and retains full activity over the course of a 24-hr reaction (data not shown), thus eliminating the possibility that the change in the rate of product formation is due to loss of active RNPs. Nuclease contamination in the protein stocks is also unlikely due to lack of DNA cleavage in the absence of sgRNA (Fig. 1B,C). Possible explanations for the change in the rate of the reaction may be that the pool of available substrate 110mer DNA1 decreases below the K_m_, SauCas9 may be subject to product inhibition and/or slow product release (Fersht, 1999), or a more complicated mechanism is responsible for this observation, which we cannot account for at this time.

**Figure 2.**
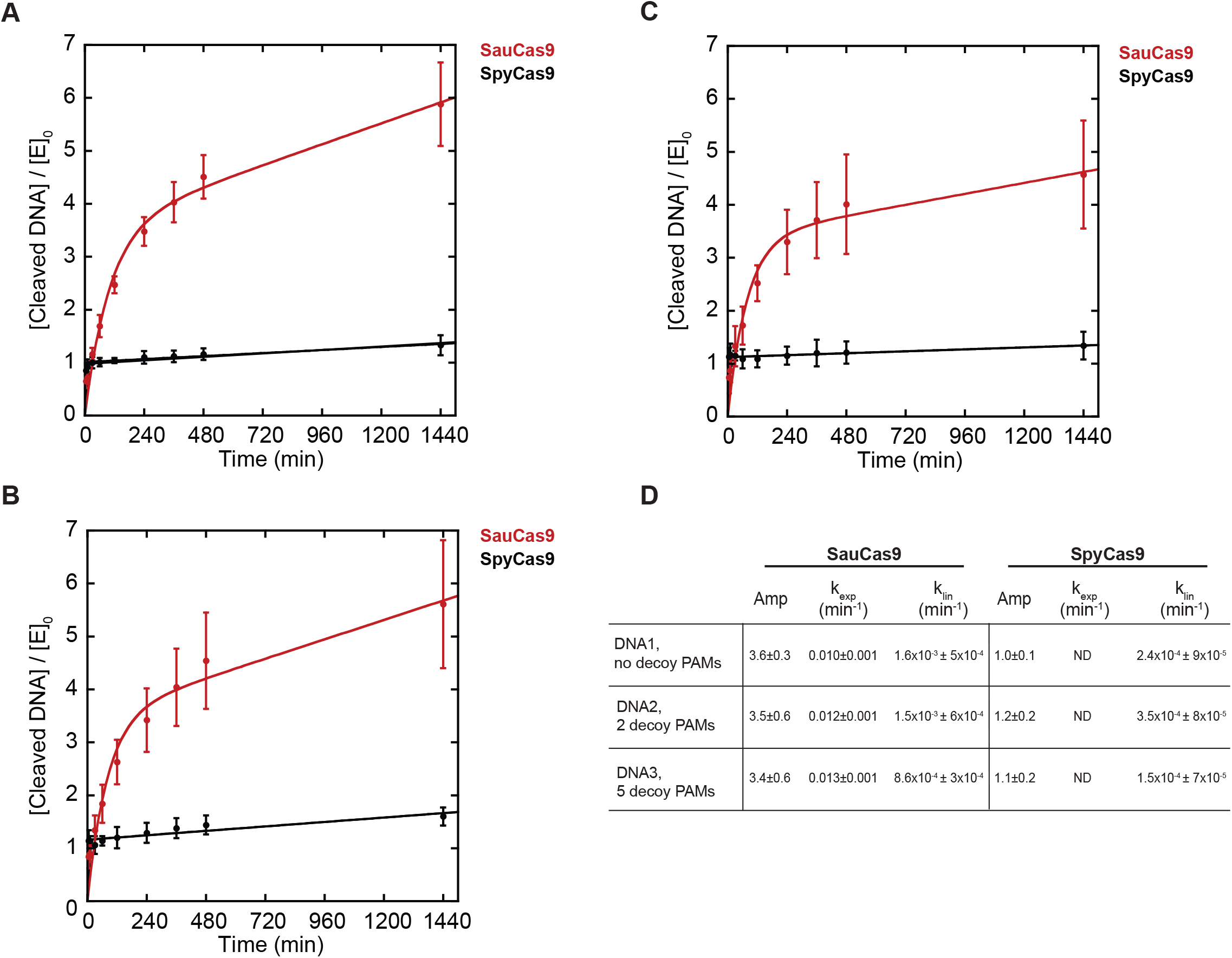
*S. aureus* Cas9 is a multiple turnover enzyme. 25 nM SpyCas9 (black) or SauCas9 (red) was pre-incubated with 100 nM sgRNA for 15 min prior to addition of 250 nM DNA. Cleaved DNA, divided by enzyme concentration, was plotted versus time. **A.** Target DNA, no decoy PAMs. SauCas9: amplitude = 3.6 ± 0.3; *k_exp_* = 0.010 ± 0.001 min^−1^; *k_lin_* = 1.6×10^−3^ ± 6×10^−4^ min^−1^. SpyCas9: amplitude = 1.0 ± 0.1; *k_lin_* = 2.4×10^−4^ ± 9×10^−5^ min^−1^. **B.** Target DNA2, 2 decoy PAMs. SauCas9: amplitude = 3.5 ± 0.60; *k_exp_* = 0.012 ± 0.001 min^−1^; *k_lin_* = 1.5×10^−3^ ± 6×10^−4^ min^−1^. SpyCas9: amplitude = 1.2 ± 0.2; *k_lin_* = 3.5×10^−4^ ± 8×10^−5^ min^−1^. **C.** Target DNA3, 5 decoy PAMs. SauCas9: amplitude = 3.4 ± 0.6; *k_exp_* = 0.013 ± 0.001 min^−1^; *k_lin_* = 8.6×10^−4^ ± 3×10^−4^ min^−1^. SpyCas9: amplitude = 1.1 ± 0.2; *k_lin_* = 1.5×10^−4^ ± 7×10^−5^ min^−1^. **D.** Summary of the data in A-C for convenience. ND – the rate of the burst could not be resolved by manual quenching. All values are reported as mean ± average deviation.

The 110mer DNA1 substrate in our study (Table 1, DNA1) contained one instance of the PAM, adjacent to the target sequence, which is not the case in genomic DNA. By chance a genome is likely to contain numerous PAMs that occur distal to the target sequence. Therefore, we tested how additional “decoy” TGGAAT PAMs – distal to the target sequence – affect the turnover activity of Spy- and SauCas9. We mutated the 110mer DNA1 substrate to contain either 2 or 5 decoy PAMs in addition to the “true” PAM located 3’- to the target sequence (Table 1). The target sequence within the DNA was not changed to avoid potential differences in editing efficiency observed with different targets and requisite sgRNAs (Doench et al., 2014; Thyme et al., 2016). Addition of extra PAMs resulted in a modest decrease in the total DNA cleaved by SauCas9, from 150 ± 20 nM in the absence of decoy PAMs to 110 ± 30 nM in the presence of 5 decoy PAMs (Fig. 2A-C). The degree of cleavage observed with SpyCas9 was not significantly affected by presence of decoy PAMs. The measurable rates of reactions were not affected significantly for either homolog (Fig. 2B-D).

In summary, SpyCas9 cleaved approximately a stoichiometric amount of DNA while SauCas9 cleaved a 6-fold excess over 24 hr, suggesting a significantly faster rate of turnover in SauCas9 than in SpyCas9, *in vitro.* The presence of decoy PAMs throughout the DNA modestly decreased the amount of cleaved product for SauCas9 but not SpyCas9.

### *S. aureus* Cas9 does not have detectable post-cleavage trimming activity

Recent work demonstrated that SpyCas9 exhibits RuvC-catalyzed post-cleavage exonuclease activity on the cleaved DNA products (Stephenson et al., 2018). CE traces analyzing DNA cleavage by SpyCas9 over 24 hr are highly consistent with this observation. The non-PAM strand, labeled with a 5’-ROX fluorophore, resulted in a single homogenous peak, while the PAM-containing strand – labeled with a 5’-FAM fluorophore – resulted in a series of smaller peaks indicating that the PAM-distal fragment of the PAM-containing strand was cut in various locations, degraded further upon cleavage or both (Fig. 3A). Surprisingly, no significant strand degradation was observed in the reaction containing SauCas9 (Fig. 3B). Both the PAM-containing (5’-FAM, cleaved by RuvC) and the non-PAM (5’-ROX, cleaved by HNH) strands yielded single homogenous peaks that increased in magnitude over the course of the reaction (Fig. 3B). Because the reaction substrates were labeled on the 5’-ends (Table 1), we cannot rule out degradation of the PAM-proximal PAM-containing strand nor the PAM-distal fragment of the non-PAM strand; however, these data provide an insight into another important mechanistic difference between the *S. pyogenes* and *S. aureus* Cas9 homologs that is to be considered in applications of the CRISPR/Cas9 technology.

**Figure 3.**
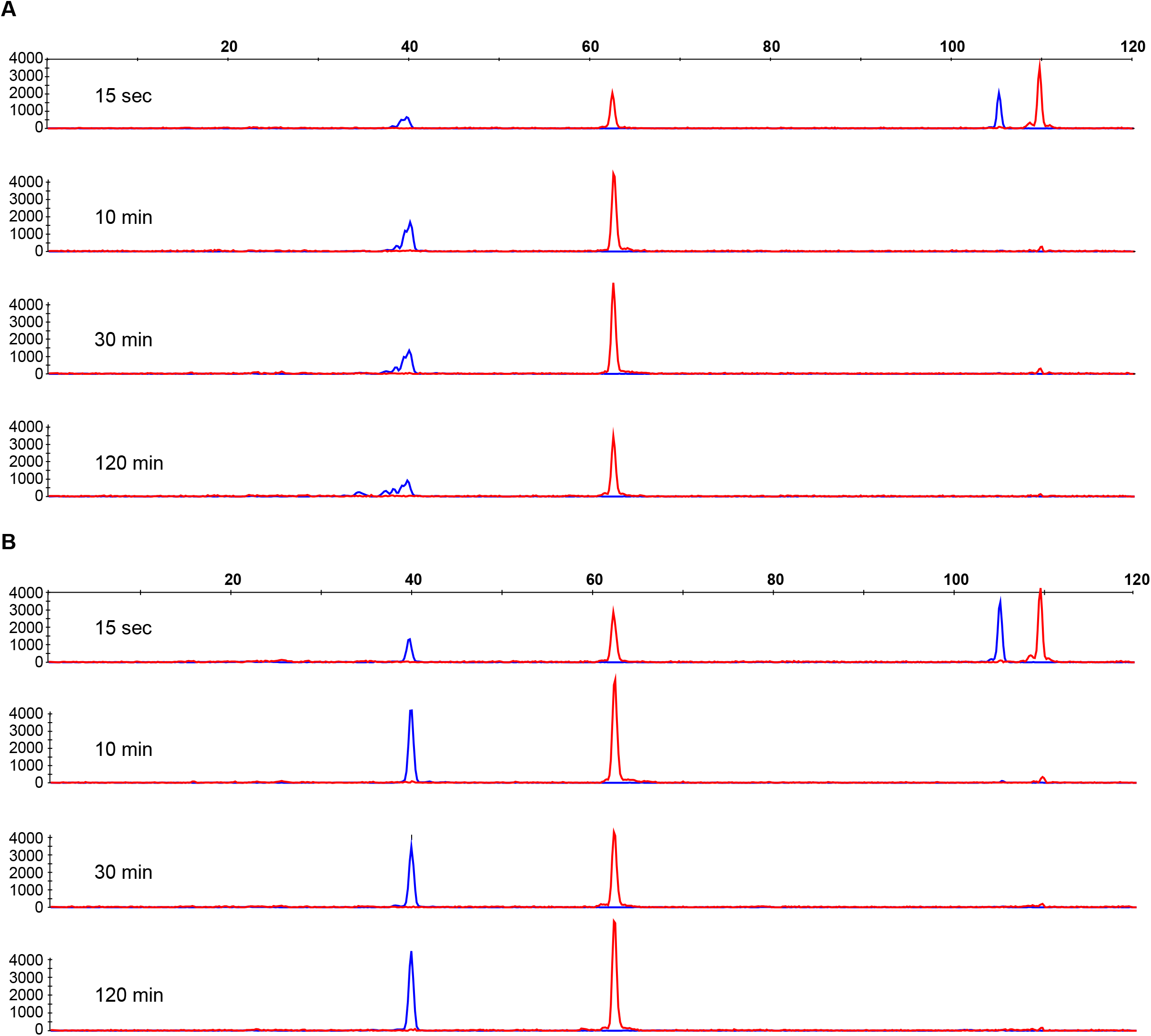
*S. aureus* Cas9 does not exhibit detectable post-cleavage trimming activity on cleaved DNA. **A–B**. Representative capillary electrophoresis of 110mer DNA labeled with 5’-FAM (blue trace) on the PAM-containing strand and 5’-ROX (red trace) on the non-PAM strand hydrolyzed by (A) SpyCas9 or (B) SauCas9 at reaction time points of 15 sec, 10 min, 30 min, and 120 min. Data are plotted as relative fluorescence versus DNA oligomer length.

## DISCUSSION

SpyCas9 is the best-characterized Cas9 enzyme due in large part to its widespread adoption as a genome editing tool. SauCas9 shares 17% sequence identity with SpyCas9 and has been less intensively reported on. Our *in vitro* data suggests that SauCas9 is a multiple-turnover enzyme while SpyCas9 is not. This finding may provide some insight into previous observations suggesting that SauCas9 is modestly more active than SpyCas9 in cells (Xie et al., 2018). DNA cleavage *in vitro* may not completely correlate with editing *in vivo;* however, the fact that SauCas9, but not SpyCas9, is able to undergo multiple rounds of catalysis, suggests a fundamental mechanistic difference between the homologs. To our knowledge, this is the first report of a multiple-turnover Cas9. The rate of turnover is slow but it is significantly faster than for SpyCas9. It is tempting to speculate that a multiple-turnover Cas9 could be required in a lower dose (e.g., for genome editing) than a Cas9 that remains bound to a cleaved target.

It is somewhat surprising that in spite of faster turnover and a longer (i.e., less common) PAM, the amount of product formation was modestly decreased in DNA2 and DNA3 – containing additional PAMs – for SauCas9 but not SpyCas9. One possible model, which will be the subject of future studies, is that while SauCas9 may have a faster rate of product release, it might bind a substrate PAM sequence with a higher affinity than SpyCas9, irrespective of the adjacent target sequence.

Recent mechanistic work demonstrates that SpyCas9, cleaves the PAM-containing strand of DNA in variable locations and that cleaved DNA is modestly degraded post-cleavage (Stephenson et al., 2018) while our data show that SauCas9 cleaves in a single location and no detectable degradation products accumulate. Stephenson *et al.* also suggest that the PAM-containing strand of the DNA is more flexible in the SpyCas9:sgRNA:target DNA complex, which might contribute to heterogeneity in the cleavage sties. In our experimental setup, CE provides single nucleotide resolution and the presence of uniform peaks for SauCas9-catalyzed reactions strongly suggests that the DNA is cleaved in a single location. It is possible that the PAM-containing strand is more rigid in the SauCas9:sgRNA:target DNA complex than in the analogous complex with SpyCas9. An alternative model consistent with our data is that SauCas9 exhibits a faster rate of product release in which SauCas9 does not have sufficient time to degrade the DNA.

In this study, we compared DNA cleavage activity of *S. pyogenes* and *S. aureus* Cas9 in the presence of saturating sgRNA, *in vitro.* Our data provides novel insights into the mechanism of catalysis for these enzymes. SauCas9 is smaller by more than 300 amino acids, greatly reducing the challenges of vector-based delivery into cells (Ran et al., 2015; Senis et al., 2014), is a multiple-turnover enzyme, and cleaves DNA in a single location without further degradation. Taken together, these findings reinforce SauCas9 as an attractive alternative to SpyCas9 for future biotechnological and therapeutic applications.

## MATERIALS AND METHODS

### Reagents

*S. pyogenes* Cas9 (# M0386M), the EnGen sgRNA Synthesis Kit, *S. pyogenes* (# E3322S), T4 RNA Ligase 1 (# M0437M), HiScribe T7 High Yield RNA Synthesis Kit (E2040S), 2x RNA Loading Dye (# B0363S), Murine RNase Inhibitor (# M0314L), Q5 Hot Start High-Fidelity 2X Master Mix (# M0494L), Monarch PCR & DNA Cleanup Kit (# T1030L), Proteinase K (# P8107S), and NEBuffer 3.1 (# B7203S) with a 1X composition of 100 mM NaCl, 50 mM Tris-HCl, 10 mM MgCl2, 100 μg/ml BSA, pH 7.9 at 25°C were all from New England Biolabs (Ipswich, MA). Both *S. pyogenes* and *S. aureus* Cas9 were purified at New England Biolabs (Ipswich, MA) using standard liquid chromatography protein purification techniques. Protein stock concentration for both Spy- and SauCas9 was measured by absorbance of 280 nm light on a NanoDrop instrument (A_280_) as well as Bio-Rad Bradford assays per manufacturer protocol. All DNA oligomers were ordered from Integrated DNA Technologies (Coralville, IA). Cytidine-5’-phosphate, containing a Cytidine-5 fluorophore (# NU-1706-CY5), used for labeling sgRNAs was ordered from Jena Bioscience (Germany). The Zymo RNA Clean & Concentrator −5 kit (#R1016) was purchased from Zymo Research (Irvine, CA). The SequaGel – UreaGel (# EC-833) system was from National Diagnostics (Atlanta, GA).

### sgRNA transcription, purification, and labeling

*S. aureus* sgRNA was transcribed using the single-stranded DNA template 5’-ATCTCGCCAACAAGTTGACGAGATAAACACGGCATTTTGCCTTGTTTTAGTAGATTCTGTTTCCAGAGTACTAAAACACAGTTCGATTAACTTTCACTATAGTGAGTCGTATTAATTTCGA and an oligo that is complementary to the T7 promoter region 5’-TCGAAATTAATACGACTCACTATAG. Oligos were transcribed using the NEB HiScribe T7 High Yield RNA Synthesis Kit according to manufacturer protocol. *S. pyogenes* sgRNA was transcribed with the EnGen sgRNA Synthesis Kit according to manufacturer protocol with the following oligo added to the reaction 5’-TTCTAATACGACTCACTATAGTGAAAGTTAATCGAACTGTGTTTTAGAGCTAGA. Transcription products were purified as described previously (Linpinsel and Conn, 2012) with small modifications: reactions were quenched with an equal volume of 2x NEB RNA Loading Dye and resolved on a 10% SequaGel (10% acrylamide, 7.5 M urea) in 1x TBE buffer. RNA bands were visualized by UV shadowing, cut out, crushed with a micro spatula, and soaked overnight at 4°C in 300 mM NaOAc, pH 5.5. Eluted RNA was filtered with 0.22 μm pore syringe-driven PVDF filter, precipitated with 3 volumes of 95% ethanol at −20°C for 16 hr, and resuspended in water. sgRNAs were labeled with 3’-Cy5 using T4 Ligase 1 in a 20 μL reaction containing 1x T4 Reaction Buffer, 2 μM RNA, 150 μM ATP, 10% DMSO, 5 μM pCp-Cy5, 4 units of murine RNase inhibitor, and 60 units of T4 RNA Ligase. Reactions were incubated at 25°C, for 2 hrs. Labeled sgRNAs were purified using the Zymo RNA Clean & Concentrator −5 kit according to manufacturer protocol.

### sgRNA binding measured by fluorescence anisotropy

sgRNA binding to Cas9 proteins was measured in NEBuffer 3.1 in a 600 μL quartz cuvette on a Horiba Scientific FluoroMax-4 fluorometer. Unlabeled Cas9 was titrated in the presence of 10 nM 3’-Cy5-labeled sgRNA. Measured changes in fluorescence anisotropy were converted to fraction bound. The data were fit with a quadratic equation as described previously (Pollard, 2010):

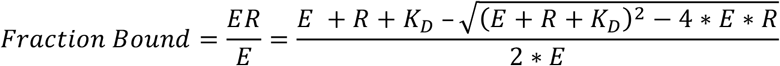

where *ER* is sgRNA bound to Cas9, *E* is the total enzyme in the reaction, and *R* is equal to the total sgRNA in the reaction. The data were fit with KaleidaGraph software.

### DNA amplification and purification

Forward and reverse DNA primers labeled with a 5’-FAM and a 5’-ROX fluorophore, respectively, were used to amplify single-stranded unlabeled oligonucleotides with the target sequence and a PAM (Supplement 1) – using the NEB Q5 Hot Start High-Fidelity 2X Master Mix per manufacturer protocol – thus creating various 110mers used in the study (Table 1). PCR reactions were purified with the Monarch PCR & DNA Cleanup Kit per manufacturer protocol.

### DNA cleavage assays

All reactions were performed in NEBuffer 3.1 (see “Reagents” above for composition) and carried out at 22°C. Single turnover experiments to determine the *k_max_* and K_1/2_ for sgRNA: 25 nM Cas9 was preincubated with different concentrations of sgRNAs for 15 min and reactions were initiated by addition of 110mer DNA (10 nM final concentration). Reaction time points were acquired by quenching 2 μL aliquots by combining with 2 μL of 2x quench mix containing 100 mM EDTA, 2% SDS, and 0.2 units/μL of Proteinase K. The quench mix was used not more than 10 min after addition of Proteinase K to insure maximum proteolytic activity. Quenched reactions were resolved on the Applied Biosystems 3730xl instrument and analyzed using the PeakScanner software (Applied Biosystems) (Greenough et al., 2016). Fraction of DNA cleaved was calculated by dividing the integrated peak area of the product peak by the combined integrated area of the product and the substrate peaks. Fraction of DNA cleaved was plotted versus time and fit with a single exponential equation:

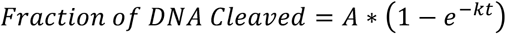

where *t* is time, *A* is amplitude and *k* is the observed rate constant, *k_obs_*.

Observed rate constants were plotted versus sgRNA concentration and fit to a hyperbolic equation:

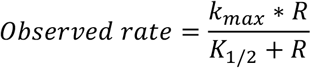

where the *k_max_* is the maximal rate of the reaction when not limited by sgRNA concentration, *R* is the concentration of sgRNA in the reaction, and *K_1/2_* is the concentration of sgRNA required to achieve half-maximal rate of DNA cleavage.

Multiple turnover reactions were performed exactly as described above except the concentration of the DNA was 250 nM. The fraction of DNA product cleaved was multiplied by 250 nM total DNA in the reaction to obtain nM product*min^−1^ and divided by 25 nM enzyme concentration to obtain rates in units of min^−1^. The reactions were fit to:

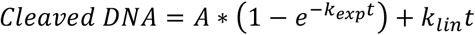

where *t* is time. *k_exp_* and *k_ln_* are the exponential and linear rates describing DNA cleavage, respectively.

## SUPPLEMENTAL MATERIAL

A complete table of all of the DNA oligomers used for *in vitro* transcription of sgRNAs and PCR amplification of DNA substrates is available online.

## ACKNOWLEDGEMENTS

We would like to thank Katherine Marks for purification of SauCas9, Siu-Hong Chan, Bill Jack, Greg Lohman, Lana Saleh, and NEB Enzymology as well as Postdoctoral clubs for productive discussions and insightful suggestions. We are also grateful to the NEB sequencing core as well as all of the facilities personnel for the analysis of capillary electrophoresis samples and daily support of science experiments.

